# Klf4 methylated by Prmt1 is required for lineage segregation of epiblast and primitive endoderm

**DOI:** 10.1101/2020.04.24.059055

**Authors:** Zhen-yu Zuo, Guang-hui Yang, Hai-yu Wang, Yan-jun Zhang, Yun Cai, Fei Chen, Yi Xiao, Mo-bin Cheng, Yue Huang, Ye Zhang

## Abstract

The second cell fate decision in the early stage of mammalian embryonic development is pivotal; however, the underlying molecular mechanism is largely unexplored. Here, we report that Prmt1 acts as an important regulator in primitive endoderm (PrE) formation. First, an embryonic chimeric assay showed that Prmt1 inhibition induces the integration of mouse embryonic stem cells (ESCs) into the PrE. Second, Prmt1 inhibition promotes Gata6 expression in both mouse blastocysts and ESCs. Single-cell RNA sequencing and flow cytometry assays demonstrated that Prmt1 depletion in ESCs contributes to an emerging cluster, where PrE genes are upregulated significantly. Furthermore, the efficiency of extraembryonic endoderm stem cell induction increased in Prmt1-depleted ESCs. Finally, we showed that the pluripotency factor Klf4 methylated at Arg396 by Prmt1 is required for recruitment of the repressive mSin3a/HDAC complex to silence PrE genes. Therefore, we reveal a regulatory mechanism for cell fate decisions centered on Prmt1-mediated Klf4 methylation.

## Introduction

Mammalian embryogenesis occurs through an ordered process (Shahbazi & Zernicka-Goetz, 2018; Zernicka-Goetz *et al*, 2009). During mouse early embryo development, the first cell fate decision results in the emergence of two cell lineages, the inner cell mass (ICM) and trophectoderm (TE) (Johnson & Ziomek, 1981; Saiz & Plusa, 2013), which are specified at the molecular level by Oct4/Pou5f1 and Cdx2, respectively (Nichols *et al*, 1998; Niwa *et al*, 2005; Strumpf *et al*, 2005). Subsequently, ICM segregates into the epiblast (EPI) and primitive endoderm (PrE) at E4.5, which is termed the second cell fate decision (Chazaud *et al*, 2006; Shahbazi & Zernicka-Goetz, 2018). The process of PrE and EPI specification within the ICM is beginning to be elucidated. To be more specific, Gata6 and Nanog, the key factors for ICM segregation, become heterogeneously expressed at the early blastocyst stage (Guo *et al*, 2010; Plusa *et al*, 2008; Xenopoulos *et al*, 2015). Subsequently, these factors progressively become mutually exclusive in their expression, which marks the initiation of lineage segregation (Plusa *et al*., 2008). Embryonic stem (ES) cells with high Nanog expression and low Gata6 expression acquire an EPI fate (Mitsui *et al*, 2003; Nakai-Futatsugi & Niwa, 2015; Xenopoulos *et al*., 2015), while PrE lineage commitment requires the repression of Nanog (Frankenberg *et al*, 2011) and sequential activation of Gata6 (Fujikura *et al*, 2002; Koutsourakis *et al*, 1999; Schrode *et al*, 2014), Sox17 (Niakan *et al*, 2010), Gata4 (Soudais *et al*, 1995) and Sox7 (Artus *et al*, 2011). Overall, Gata6 and Nanog, the earliest markers of the PrE and EPI lineages, respectively, act near the top of the hierarchy regulating PrE development (Frankenberg *et al*., 2011). However, the molecular mechanisms governing the specification of PrE and EPI, or rather regulating the switch of Gata6 and Nanog expression, are still poorly understood.

Epigenetic modifications have been proven to be essential for cell fate determination and early embryo development (Atlasi & Stunnenberg, 2017; Jambhekar *et al*, 2019). Protein arginine methylation catalyzed by enzymes in the protein arginine methyltransferase (PRMT) family is a prevailing posttranslational modification that functions as an epigenetic regulator of transcription in multiple cellular processes, including cell fate decisions (Blanc & Richard, 2017; Guccione & Richard, 2019; Vougiouklakis *et al*, 2017; Yang & Bedford, 2013). In the PRMTs family, Prmt4/Carm1 regulates cell fate decisions early in the four-cell stage by methylating histone H3R26 and activating Sox21 expression in mouse ESCs (Goolam *et al*, 2016; Torres-Padilla *et al*, 2007). We previously reported that Sox2 is methylated by Carm1 at R113, leading to its self-association and activation (Zhao *et al*, 2011). Prmt5 can methylate histone H2A to repress differentiation genes in ES cells and is crucial for early mouse development (Tee *et al*, 2010). Prmt6 regulates both pluripotency genes and differentiation markers to participate in the maintenance of ES cell identity (Lee *et al*, 2012). Although Prmt1 provides the bulk of the protein arginine methylation activity within cells (Bedford & Clarke, 2009; Simandi *et al*, 2015) and Prmt1-null mice die at E6.5 (Pawlak *et al*, 2000), little is known about the role of Prmt1 in early embryonic development.

In this study, we report that Prmt1 depletion in mouse ESCs results in fluctuating heterogeneity and leads to the emergence of a cluster with the properties of PrE progenitor. Prmt1 interacts with and arginine-methylates Klf4, a key pluripotency factor. Methylated Klf4 at Arg396 is essential for the recruitment of the repressive mSin3a/HDAC complex to silence the expression of PrE genes in mouse ESCs. Both Klf4 R396 mutation and the enzymatic inactivation of Prmt1 derepress PrE genes, followed by a tendency toward the development of extraembryonic endoderm stem (XEN) cells. Moreover, a Prmt1 enzymatic inhibitor promotes the formation of PrE *in vivo*. Our data reveal a novel molecular mechanism controlling second cell fate determination in embryogenesis mediated by the enzymatic functions of Prmt1.

## Results

### Prmt1 regulates the PrE commitment *in vivo*

To investigate the function of Prmt1 in mouse embryo development, the Prmt1-specific inhibitor furamidine dihydrochloride (Fur) was employed to treat mouse AB2.2 ES cells (Huang *et al*, 2012) stably expression two different fluorescent proteins GFP and td-Tomato transgene. Then the treated ESCs were injected into the mouse E3.5 blastocyst and E6.5 chimeric embryos were dissected and observed with a fluorescence stereomicroscope (FSM) and confocal laser stereomicroscope (CLSM) (Figure 1A). We found that Fur-treated GFP^+^/td-Tomato^+^ (Fur +) cells were able to integrate into both EPI and PrE, while GFP^+^/td-Tomato^+^ cells were integrated into only EPI in the control group (Fur -) (Figure 1B). Further observations of chimeric embryos at E12.5 and mice after birth showed that the Fur-treated ES cells were still capable of integration (Figure S1). The results suggested that Prmt1 is likely to participate in the second cell fate decision during mouse embryo development, potentially by restricting the PrE commitment. To confirm the function of Prmt1 in early embryogenesis, mouse blastocysts were treated with Fur. Immunofluorescence (IF) images showed that Gata6, a marker of the PrE, was induced, while the level of Nanog, a marker of EPI, was reduced by Fur treatment in mouse blastocysts (Figure 1C). Inhibition of Prmt1 significantly increased the percentage of Gata6-positive cells (Figure 1D). Furthermore, IF showed that Fur treatment induced Gata6 gene expression while reducing the expression of Nanog in cultured E14 ESCs (Figure 1E). Moreover, RT-qPCR showed that inhibition of Prmt1 induced the expression of the *Gata4* and *Gata6* genes at the mRNA level (Figure 1F). Taken together, our data indicate that inhibition of Prmt1 can induce the expression of Gata6 to bias the fate of pluripotent cells toward the PrE lineage.

**Figure 1.**
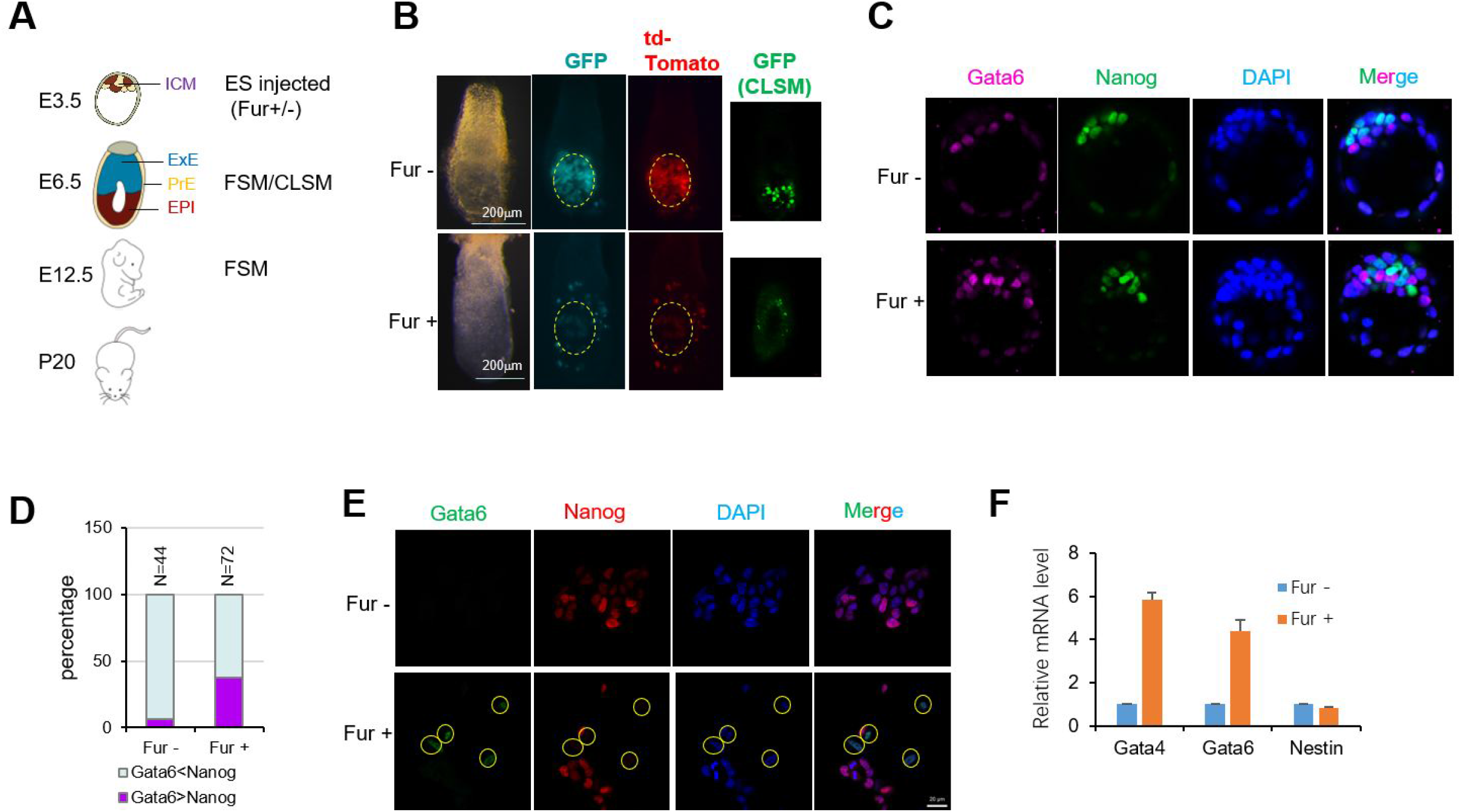
Prmt1 inhibition affects PrE formation. (A) Scheme of the chimeric assay. (B) Chimeric assay of ES cells with (Fur+) or without (Fur-) treatment. The ES cell line was AB2.2 cells with GFP and td-Tomato. After Fur treatment for 24 hr, cells were microinjected into blastocysts. Embryos were observed at E6.5. GFP (green) and Td-Tomato (red) signals were detected. Dashed lines mark epiblasts. (C) IF assays showed the effect of Fur on mouse blastocysts. Blastocysts were cultured with (Fur+) or without (Fur-) for 24 hr and then immunostained with Gata6 (red) to indicate PrE and Nanog (green) to indicate EPI. (D) Distribution of two cell types in the ICM. (E) IF assays showed the impact of Fur on the expression of Gata6 (green) and Nanog (red) in E14 cells with XEN induction. Nuclei were visualized with DAPI staining (blue). Yellow circles indicate Gata6-positive cells. (F) Fur treatment induced the expression of Gata4 and Gata6 genes using RT-qPCR, and Nestin was used as a negative control.

### Prmt1 depletion induces PrE gene expression in mouse ESCs

To illustrate the effects of Prmt1 in mouse ES cells, *Prmt1* knockout E14 cell lines (KO-1 and KO-2) were generated by using CRISPR-Cas9 technology (Cai *et al*, 2015). Depletion of Prmt1 was confirmed by western blotting (Figure 2A). The expression of Prmt1 mRNA was also significantly decreased in *Prmt1*-KO1 and *Prmt1*-KO2 cells compared with WT cells (Figure S2A and S2B). In addition, disruption of the Prmt1 genomic sequence within the knockout cell lines was confirmed by PCR and Sanger sequencing, which proved the success of Prmt1 gene editing (Figure S2C). To exclude the possibility of off-target effects caused by CRISPR-Cas9 technology, we amplified and sequenced the potential off-target loci to confirm the integrity of the genome outside of the Prmt1 gene locus. All the loci tested remained intact, which diminishes the possibility of the existence of off-target effects (Table S1). Western blot assays of extracted histones showed that the levels of asymmetric dimethylation of histone 4 arginine 3 (H4R3me2a), the predominant substrate of Prmt1, decreased significantly in *Prmt1* KO cells, while the levels of H3R17me2a, the main substrate of Carm1, did not change (lower panels, Figure 2A). All these results confirmed that Prmt1 was functionally depleted in the KO1 and KO2 cell lines.

**Figure 2.**
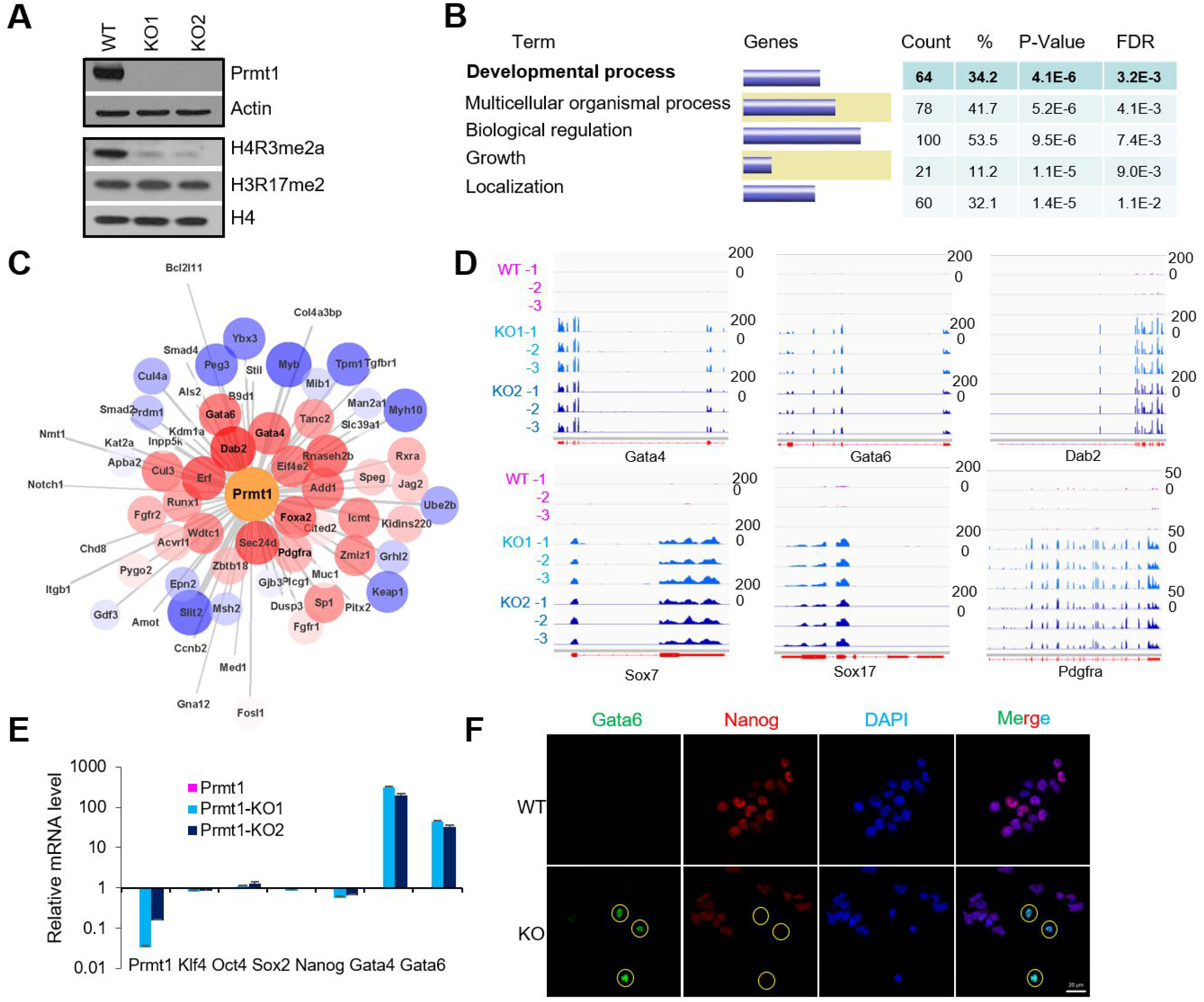
Prmt1 depletion induces PrE genes in mouse ES cells. (A) Western blotting showed the expression of Prmt1 and its impact on the levels of arginine-methylated histones (H4R3me2a and H3R17me2) in WT, Prmt1 KO1, and KO2 cells. Histone H4 was used as an internal control. (B) Enriched GO terms for Prmt1 -affected genes according to the bulk RNA-Seq data of WT and Prmt1 KO cells. (C) A WGCNA network was constructed based on Prmt1 and its related embryonic genes. Red indicates upregulation, and blue indicates downregulation. (D) Integrative Genomics Viewer (IGV) browser view of RNA-Seq reads covering the indicated genes in WT and Prmt1 KO cells. The x-axis indicates the genomic location, and the y-axis represents the normalized scale of RNA-Seq reads. (E) Expression of the indicated genes using RT-qPCR. (F) IF assays of endogenous Gata6 (green) and Nanog (red) in WT and Prmt1 KO cells. Nuclei were visualized with DAPI staining (blue). Yellow circles indicate Gata6-positive cells.

To investigate the impact of Prmt1 depletion in E14 cells, the stemness of mouse ES cells was evaluated. Alkaline phosphatase (AP) staining assays showed that *Prmt1* KO ES cells could still form distinct AP-positive colonies, although the typical tightly-packed colony morphology cannot be observed in KO cells (Figure S3A). In addition, teratoma formation assays were performed to characterize the pluripotency of *Prmt1* KO cells, and the results showed that *Prmt1* KO cells could form teratomas with differentiated structures of three germinal layers (Figure S3B). These results indicated that ES cells maintained comparable differentiation potentials after depletion of Prmt1. Next, we examined the expression of pluripotency factors by RT-qPCR and western blot assays. We found that Prmt1 depletion did not substantially change the expression of the Oct4, Sox2, and Klf4 genes at the protein and mRNA levels, while a reduction in Nanog was observed in Prmt1 KO ES cells (Figure S3C-E). Taken together, these data suggested that Prmt1 depletion did not significantly decrease pluripotency or ES cell stemness.

To further investigate the effects of Prmt1 depletion, RNA-Seq using total RNA extracted from WT and *Prmt1* KO cells was performed. Differential gene expression analysis showed that 2363 genes were downregulated after Prmt1 depletion compared to WT cells, while 2499 genes were upregulated (Figure S3E). GO analysis according to the differentially expressed genes (DEGs) revealed significant enrichment of genes involved in developmental processes (Figure 2B). To understand what developmental procedures may be affected by *Prmt1* knockout, we filtered out all the genes that have been reported to affect development from all the DEGs and analyzed their correlation with Prmt1 depletion. To our surprise, weighted gene coexpression network analysis (WGCNA) (Langfelder & Horvath, 2008) showed that upregulation of PrE genes, including *Gata6, Gata4, Dab2, Foxa2*, and *Pdgfra*, exhibited the closest relationship with Prmt1 depletion, which indicated that Prmt1 depletion could be involved in PrE formation by regulating PrE genes (Figure 2C). Significant upregulation of PrE genes in *Prmt1* KO cells could be identified from the RNA-Seq data (Figure 2D). Using qRT-PCR assays, we confirmed that the expression of *Gata4* and *Gata6* was significantly increased in *Prmt1* KO cells compared to WT cells (Figure 2E). Consistent with previous results, no significant changes were observed in the RNA levels of the pluripotency factors *Oct4, Sox2* and *Klf4* in the RNA-Seq data, while *Nanog* expression showed a minor reduction (Figure S3F). Furthermore, IF staining with antibodies specific for Gata6 (green) and Nanog (red) in WT and *Prmt1* KO cells showed that endogenous Gata6 was induced in some *Prmt1* KO cells, while, conversely, the Nanog level was reduced (Figure 2F). Interestingly, Gata6 was heterogeneously expressed in *Prmt1* KO cells, which suggested that Prmt1 might be a regulatory factor for the heterogeneity of mouse ES cells. Taken together, our data indicated that *Prmt1* knockout induced the expression of PrE genes but did not significantly affect the pluripotency of ES cells.

### Prmt1 depletion leads to the emergence of progenitors of PrE

To further investigate the impact of Prmt1 on the regulation of heterogeneity in mouse ES cells, single-cell RNA sequencing (scRNA-Seq) was performed in WT and *Prmt1* KO cells using the Chromium system (10x Genomics) (Klein *et al*, 2015; Macosko *et al*, 2015). scRNA-Seq data were obtained from 13824 individual cells, including 5955 WT cells and 7869 *Prmt1* KO cells. Nonlinear dimensionality reduction was performed using *t*-distributed stochastic neighbor embedding (tSNE) by Seurat (*K*-means=8), a visual analytics tool for integrated analysis, and 8 clusters (1-8) were identified in the integrated data from WT and KO cells (Figure 3A). Among these clusters, clusters 1 and 2 were composed of only KO cells, while clusters 3-8 contained both WT and KO cells (Figure 3B). The gene expression profiling data for all single cells allowed us to deconstruct the population heterogeneity. Principal component analysis (PCA) based on pseudotime showed that cluster 1 was distant from the other clusters (Figure S4A). The top 2000 DEGs were analyzed, and the results showed that cluster 1 was marked by upregulated expression of PrE marker genes, including *Dab2, Gata6, Foxq1, Sox17, Gata4, Foxa2*, and *Pdgfra* (Figure 3C-E and Figure S4B). Interestingly, the expression of pluripotency genes *Oct4/Pou5f1, Klf4, Sall4* and *Sox2*, but not *Nanog*, were maintained in Prmt1 KO cells (Figure 3E and Figure S3F). Combined with data from the above RNA-Seq and immunostaining experiments (Figure 2F), the gene expression profile strongly suggested that the unique emerging cluster 1 from *Prmt1* KO cells (cluster 1) was the progenitor of PrE cells. To gain insights into the gene expression dynamics and trajectory of this cluster, RNA velocity analysis was performed to predict gene expression changes in single cells (La Manno *et al*, 2018; Plass *et al*, 2018; Svensson & Pachter, 2018). The results showed a strong directional flow toward progenitors of PrE cells (Figure 3F). These results indicated that Prmt1 depletion led to the emergence of a cell population with features of progenitors of PrE within mouse ES cells.

**Figure 3.**
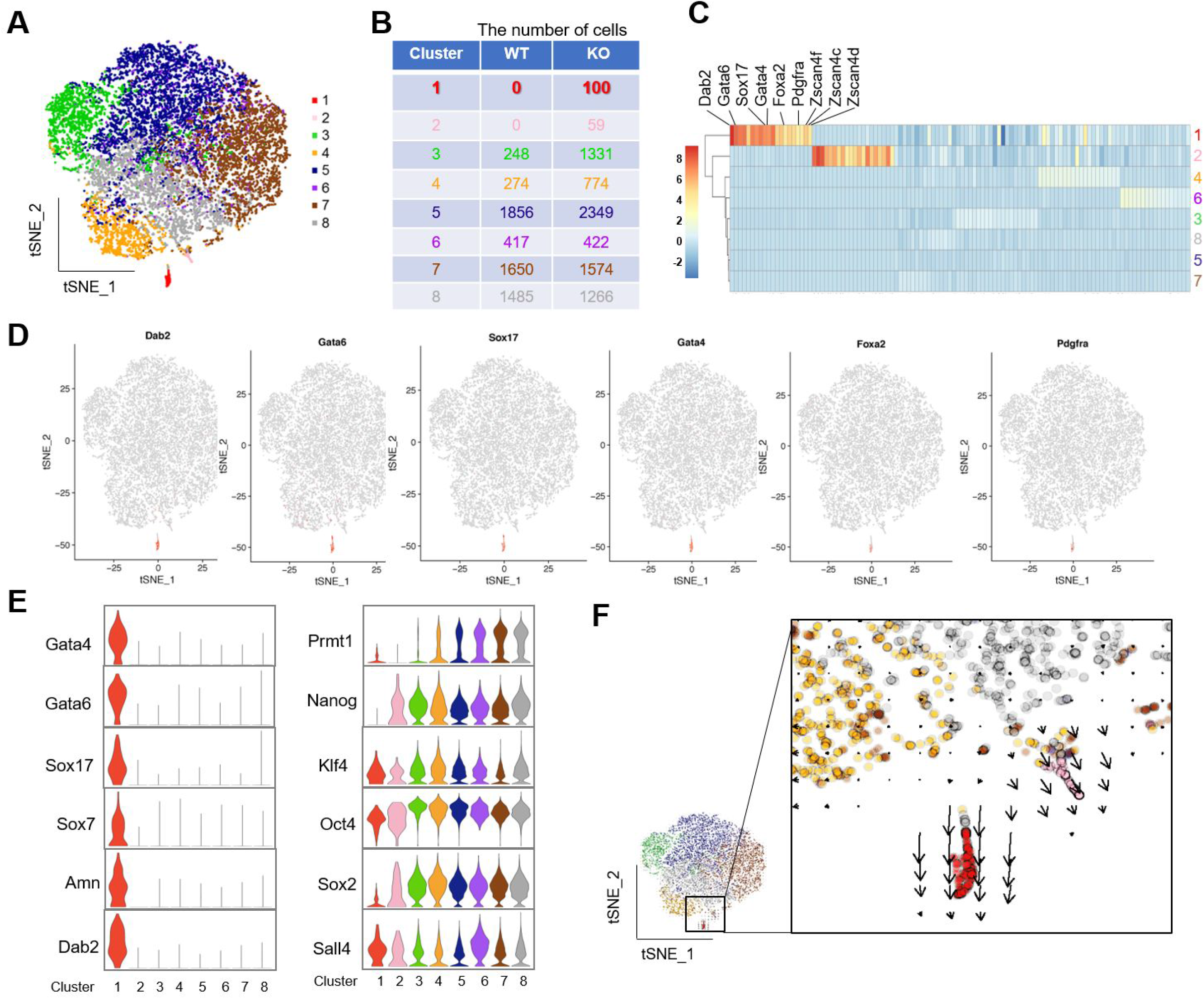
ScRNA-Seq analysis of Prmt1-depleted cells. (A, B) tSNE projection of 13,805 integrated cells, including 5930 WT cells and 7875 Prmt1 KO cells. Eight transcriptionally distinct clusters were identified (clusters 1-8) by unsupervised classification (A). Number and origin of cells in each cluster (B). (C) Heatmap displaying the expression of marker genes in each cluster. (D) Cluster 1 is marked with PrE genes indicated. (E) PrE genes showed significantly and specifically high expression in cluster 1. (F) RNA velocity of single cells shows the transcription rate (arrows).

### Prmt1 decreases the differentiation potential of mouse ES cells toward an XEN fate

Since Prmt1 depletion leads to the emergence of progenitors of PrE, we hypothesized that Prmt1 is a safeguard of mouse ES cells, preventing the cells from differentiating toward a PrE fate. XEN cells represent an *in vitro* variant of stem cells representative of the extraembryonic endoderm fate (Debeb *et al*, 2009; Kunath *et al*, 2005). To characterize how Prmt1 induces XEN fate, chemical induction of mouse E14 cells was performed as reported previously (Niakan *et al*, 2013) to analyze the ratio of XEN-induced cells using a fluorescence-activated cell sorting (FACS) assay. The results showed that *Prmt1* KO cells had higher efficiency of XEN induction than WT cells (Figure 4A), which indicated that Prmt1 depletion made the ES cells prone to an XEN fate. Interestingly, without induction, approximately one percent of the *Prmt1* KO cells showed XEN features (Figure 4A upper panel), as represented by high expression of Gata6 and low expression of Nanog (Gata6^+^/Nanog^-^). This ratio is consistent with the number of cells within cluster 1 in the scRNA-Seq data (Figure 3A and B). Furthermore, after XEN induction, more than 30% of the *Prmt1* KO cells acquired features of XEN cells, while only 1.55% of WT cells successfully became XEN cells (Figure 4A). These results strongly supported the hypothesis that Prmt1 acts as a gatekeeper of ES cell differentiation toward XEN cells.

**Figure 4.**
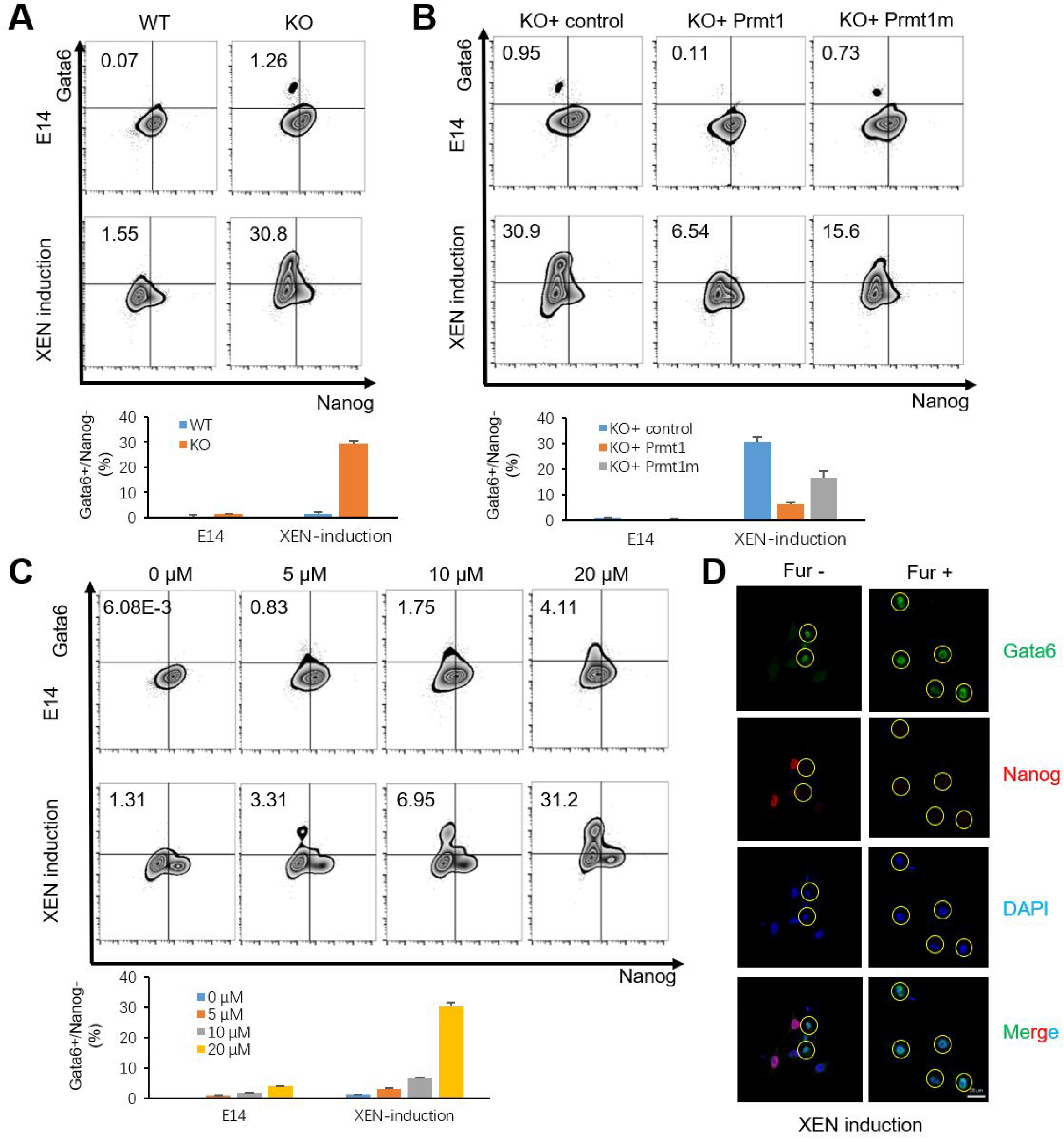
Prmt1 depletion accelerates XEN induction in mouse ES cells. (A) FACS was performed by labeling Gata6 and Nanog in WT and Prmt1 KO cells without or with XEN induction (upper panels). The percentage of Gata6+/Nanog- cells is shown in the lower panel. (B) FACS showed the impact of Prmt1 rescue in KO cells. Based on the Prmt1 KO background, a Prmt1 rescue (Prmt1) and an enzyme-inactive (Prmt1m) cell line were generated and used for XEN induction (upper panels). The percentage of Gata6+/Nanog- cells is shown in the lower panel. (C) FACS showed the impact of an inhibitor of Prmt1 (Fur) in E14 cells with and without XEN induction (upper panels). The percentage of Gata6+/Nanog- cells is shown in the lower panel. (D) IF assays showed the impact of Fur on the expression of Gata6 (green) and Nanog (red) in E14 cells with XEN induction. Nuclei were visualized with DAPI staining (blue). Yellow circles indicate Gata6+ cells.

To ascertain the role of Prmt1 in XEN induction, Prmt1 was rescued in *Prmt1* KO cells by re-expressing Prmt1 and its catalytically inactive mutant E153Q (Prmt1m) (Zhang & Cheng, 2003). Re-expression of Prmt1, but not Prmt1m, almost completely eliminated the XEN population within *Prmt1* KO cells without induction and clearly reduced the proportion of Gata6^+^/Nanog^-^ cells (6.54% vs. 30.8%) (Figure 4B). The results confirmed that Prmt1, especially *via* its enzymatic function, decreased the differentiation potential of ES cells toward XEN cells. Furthermore, we verified that Fur treatment increased the Gata6^+^/Nanog^-^ cell proportion both with and without chemical induction in a dose-dependent manner (Figure 4C), while Fur no longer changed the expression of Prmt1 (Figure S5). Moreover, IF assays showed that Fur treatment increased the number of Gata6^+^/Nanog^-^ cells among XEN-inducing E14 cells (Figure 4D). Taken together, our results demonstrate that Prmt1, specifically its enzymatic function, could prevent spontaneous differentiation of ES cells to XEN cells.

### Klf4 is involved in Prmt1-mediated chromatin remodeling at PrE genes

To gain insight into the mechanism by which Prmt1 regulates PrE genes related to the emergence of XEN cells, DNase I sensitivity assays were performed as previously described (Cheng *et al*, 2014) to investigate whether Prmt1 could affect chromatin accessibility at the promoters of PrE genes. Our data showed that Prmt1 depletion made the chromatin of PrE genes, including the *Gata4* and *Gata6* genes, more accessible at the promoter regions (Figure 5A). ChIP-qPCR assays revealed that compared to those in WT cells, the levels of histone acetylation at H3 and H4 (H3ac and H4ac) in the promoters of the *Gata4* and *Gata6* genes were dramatically increased in *Prmt1* KO cells, while other modifications (repressive H3K9me2, K3K9me3, H3K27me3, and active H3K4me3) were not changed (Figure 5B). It must be mentioned that the levels of H4R3me2 on the *Gata4* and *Gata6* genes were low in WT cells and did not change in *Prmt*1 KO cells, although the global levels of this modification decreased in Prmt1 KO cells (middle panels, Figure 2A), which implied that the decrease in H4R3me2 might make a small contribution to the activation of target genes in the *Prmt1* KO condition. In addition, hyperacetylation of H3 and H4 mediated by Prmt1 depletion was also observed in the promoters of PrE genes, including *Sox17, Sox7, Pdgfra*, and *Amn* (Figure S6). These results indicated that altered histone acetylation rather than histone arginine (or lysine) methylation could be the cause of chromatin changes at PrE gene promoters after Prmt1 depletion. Furthermore, ChIP-qPCR assays were performed by using antibodies specific for the mSin3a complex (HDAC1, HDAC2, and mSin3a) and the heterochromatic complex (HP1). The results showed that HDAC1, HDAC2, and mSin3a, but not HP1, were recruited to the promoters of the *Gata4* and *Gata6* genes (open bars, Figure 5C). Importantly, the recruitment of the mSin3a complex was significantly reduced in Prmt1-depleted ES cells (filled bars, Figure 5C) for further activation of the *Gata4* and *Gata6* genes.

**Figure 5.**
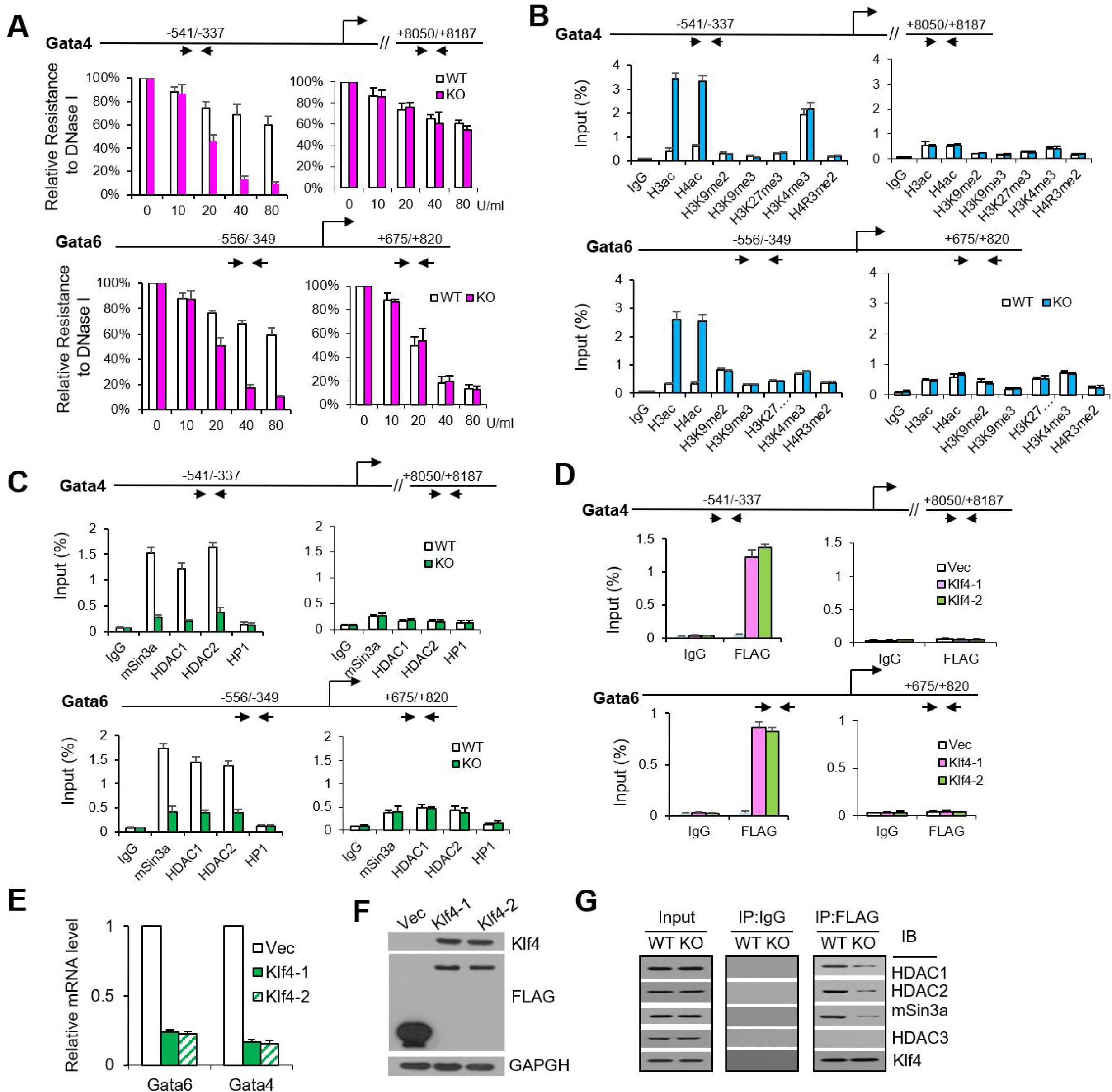
Klf4 is involved in chromatin remodeling of PrE genes mediated by Prmt1. (A) DNase I sensitivity analysis showed open chromatin in the promoter regions of the Gata4 and Gata6 genes in WT (open bars) and Prmt1 KO (filled bars) cells. The indicated amounts of DNase I (0, 10, 20, 40, and 80 U/ml) were used. (B and C) ChIP-qPCR analysis of the occupancy of histone modifications (H3ac, H4ac, H3K9me3, H3K27me3, H3K4me3, and H4R3me2) (B) and the repressive complex (mSin3a, HDAC1, HDAC2, and HP1) (C) in the upstream region of Gata4 and Gata6 in WT (open bars) and Prmt1 KO (filled bars) cells. Regions +8050 of Gata4 and +733 of Gata6 were used as negative control regions for the genes. An anti-FLAG antibody was used for ChIP. IgG was used as a ChIP control. (D) ChIP-qPCR analysis of ectopic FLAG-Klf4 binding to the promoters of Gata4 and Gata6 in E14 cells. (E and F) Stable ectopic expression of Klf4 (Klf4-1 and Klf4-2) was confirmed by western blotting (E), and RT-qPCR showed the effect of ectopic Klf4 on the expression of the Gata4 and Gata6 genes (F). Empty vector was transfected into E14 cells as a control (Vec). (G) Co-IP assays of Klf4 with the mSin3a complex in E14 cells. Whole-cell extracts (WCEs) of WT and Prmt1 KO cells were subjected to IP with an anti-Klf4 antibody and then blotted with antibodies specific for HDAC1, HDAC2, mSin3a, Klf4 and HDAC3 as a negative control. WCEs that were not subjected to IP were used as the input. IgG was used as a negative control.

Interestingly, the promoter regions of PrE genes described above contain putative Klf4-binding sites (CACCC), including the −541/−337 bp region for *Gata4* (−340 bp) and −556/−349 bp for *Gata6* (−496 bp). Klf4 belongs to the Krüppel-like family (KLF) proteins, which have been reported to be involved in many cellular processes, including pluripotency maintenance and mouse embryonic development (Aksoy *et al*, 2014; Bialkowska *et al*, 2017; Takahashi *et al*, 2007; Takahashi & Yamanaka, 2006). To address the mechanistic role of Klf4 in Prmt1-mediated gene expression, stable E14 cell lines with ectopic expression of Klf4 were generated by infection with lentiviral vectors expressing FLAG-Klf4 (designated Klf4-1 and Klf4-2), with a GFP lentiviral vector as a control. Ectopic expression of Klf4 was confirmed by western blotting (Figure 5E). ChIP-qPCR assays showed that Klf4 occupied the putative Klf4-binding sites in the promoter regions of the PrE genes *Gata4* (−340 bp) and *Gata6* (−496 bp) (Figure 5D). RT-qPCR assays showed that the mRNA levels of *Gata4* and *Gata6* were reduced by ectopic Klf4 expression in E14 cells (Figure 5F), which was consistent with a previous report (Aksoy *et al*., 2014). These results suggested that Klf4 might inhibit the expression of *Gata4* and *Gata6 via* direct recruitment to the promoters of these genes in mouse ES cells. Furthermore, Co-IP assays showed that the interaction between Klf4 and the mSin3a complex was weaker in Prmt1 KO cells than in WT cells (Figure 5G). Taken together, our data suggest that Klf4 occupancy at PrE gene promoters is involved in the inhibition of PrE gene expression *via* the mSin3a complex and that depletion of Prmt1 could alleviate this repression and lead to the activation of PrE genes in mouse ES cells.

### Klf4 is methylated by Prmt1 at arginine 396 and required for repression of PrE genes in mouse ES cells

Considering that both Prmt1 enzymatic activity and Klf4 occupancy are involved in PrE gene expression, we hypothesized that Prmt1 might methylate Klf4 directly and affect its regulatory functions. To confirm this hypothesis, we examined the interaction between Prmt1 and Klf4. A coimmunoprecipitation (Co-IP) assay was performed in HEK293T cells cotransfected with FLAG-Prmt1 and Myc-Klf4. The results showed that Prmt1 was detected with an anti-FLAG antibody after IP with an anti-Myc antibody, demonstrating an interaction with Klf4 (Figure 6A). In addition, FLAG-Klf4 from HEK293T cells was pulled down by purified GST-Prmt1 but not by GST alone (Figure 6B), which further confirmed the interaction between Prmt1 and Klf4. To identify the binding region of Klf4 that was responsible for the interaction with Prmt1, FLAG-tagged truncations of Klf4 were constructed. Pulldown assays using GST-Prmt1 indicated that the 401-483 aa region of Klf4 is responsible for its association with Prmt1 (Figure 6C). By using purified GST-Klf4 and its truncations, GST pulldown assays for FLAG-Prmt1 confirmed that the fragment from 401-483 aa of Klf4 bound to Prmt1; moreover, the region from 386-400 aa impeded the interaction between these proteins (Figure 6D). Next, *in vitro* methylation assays were performed using purified GST-Klf4 and its truncations and Prmt1 purified from HEK293T cells in the presence of ^3^H-SAM. The results showed that full-length Klf4 (1-483 aa) was methylated in the presence of Prmt1 but not Prmt1m (Prmt1 E153Q mutation) (lane 3 *vs* lane 4, Figure 6E), and GST alone was not methylated by Prmt1. Notably, ^3^H signals were also observed on the 386-400 aa and 1-400 aa fragments but not on other derivatives (lanes 9 and 11, Figure 6E), which indicated that the arginine methylation site should be located between 386 and 400 aa in Klf4. Furthermore, an *in vitro* methylation assay followed by mass spectrometric (MS) analysis showed that arginine residue R396 was dimethylated by Prmt1 (Figure 6F). No ^3^H signal was detected in the methylation assay when arginine 396 of Klf4 was mutated to lysine (R396K) (Figure 6G), which confirmed that R396 of Klf4 was methylated by Prmt1.

**Figure 6.**
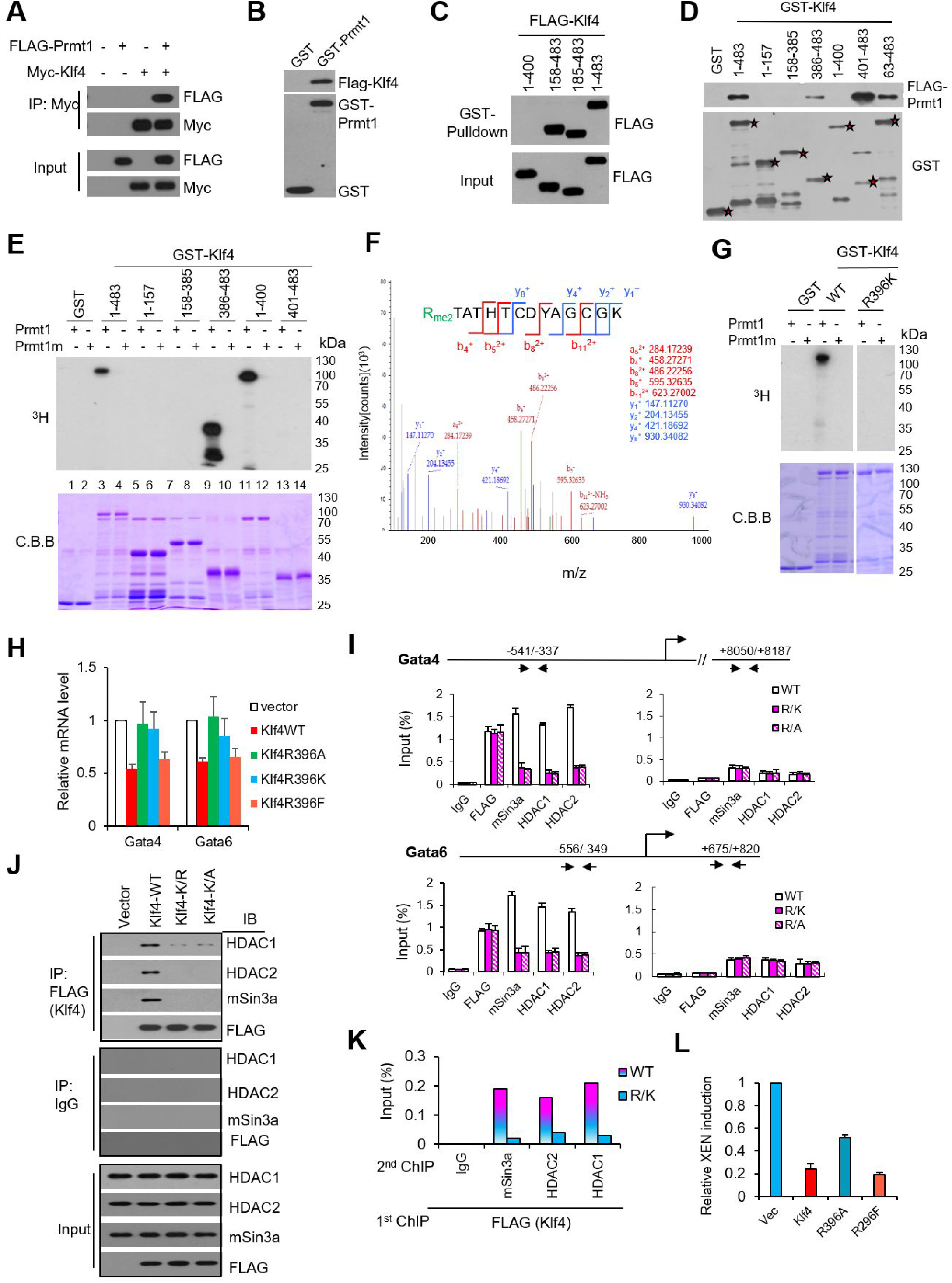
Klf4 is methylated by Prmt1 at arginine 396, and methylated Klf4 is required for Prmt1-mediated expression of PrE genes. (A) Co-IP of Prmt1 and Klf4. Whole-cell extracts (WCEs) of HEK293T cells transfected with (+) or without (-) Myc-Klf4 and/or FLAG-Prmt1 were subjected to IP with an anti-Myc antibody and blotted with antibodies specific for FLAG for Prmt1 and Myc for Klf4. (B) GST pulldown assays to detect the interaction of Prmt1 with Klf4. GST or GST-Prmt1 was incubated with WCEs of HEK293T cells expressing FLAG-Klf4 and then blotted with antibodies specific for FLAG or GST. (C) Purified GST-Prmt1 was incubated with WCEs of HEK293T cells expressing FLAG-Klf4 and its truncations. The GST pulldown products were immunoblotted with an anti-FLAG antibody, and WCEs that were not subjected to IP were used as the input. (D) Purified GST-Klf4 and its derivatives were incubated with WCEs of HEK293T cells expressing FLAG-Prmt1 or vector. The GST pulldown products were then immunoblotted with an anti-FLAG antibody for Prmt1 or an anti-GST antibody. (E) Autoradiography of in vitro methylation assays using purified GST-Klf4 and its derivatives with Prmt1 or Prmt1m (an inactive enzymatic mutant). Total amounts of GST-Klf4 and Klf4 truncations were visualized by Coomassie brilliant blue (C.B.B.) staining. (F) MS analysis of a Klf4 peptide mixture to identify methylated sites in vitro. Dimethylated arginine (Rme2) is displayed in green. (G) Autoradiography of in vitro methylation assays using purified GST-Klf4 and an R396K mutant of Klf4 with Prmt1 or Prmt1m. (H) RT-qPCR analysis showed the effects of Klf4-WT, Klf4-R396A, Klf4-R396K, Klf4-396F, and vector on the expression levels of Gata4 and Gata6 in E14 cells. (I) ChIP assays showed the recruitment of mSin3a, HDAC1, and HDAC2 to the promoters of Gata4 and Gata6 in E14 cells transfected with Klf4 (WT) or Klf4 point mutants at R396 (R/K or R/A). (J) Co-IP assays of Klf4 with the mSin3a complex. WCEs of E14 cells transfected with FLAG-Klf4 (WT) or its R396 mutants (Klf4-R/K or Klf4 R/A) were subjected to IP with an anti-FLAG antibody and then blotted with antibodies specific for HDAC1, HDAC2, mSin3a and FLAG for Klf4. WCEs that were not subjected to IP were used as the input. IgG was used as a negative control. (K) ChIP/re-ChIP assays showed that Klf4-mediated mSin3a/HDAC recruitment to the promoter of Gata6 is arginine methylation dependent. E14 cells were transfected with FLAG-Klf4 (WT) or its mutant at R396 (R/K). An anti-FLAG antibody was used for the initial ChIP (1st) to obtain the Klf4-associated chromatin fragments. Then, these fragments were subjected to re-ChIP (2nd) using mSin3a, HDAC1, and HDAC2 antibodies. IgG was used as a ChIP control. (L) FACS showed the impact of mutant R396 of Klf4 in E14 cells transfected with Klf4 and its mutants R396A and R396F. The percentage of Gata6+/Nanog- cells is shown.

To investigate the role of arginine-methylated Klf4 in regulating ES cell fate, R396 mutations were used to examine the effect of Klf4 on PrE genes. RT-qPCR assays showed that the R396A or R396K mutation of Klf4 was no longer able to suppress the expression of Gata4 and Gata6, while the R396F mutation, a mimic of methylated arginine, still suppressed Gata4 and Gata6 expression, with an effect similar to that of WT Klf4 (Figure 6H). ChIP-qPCR data showed that the mSin3a complex (mSin3a, HDAC1, and HDAC2) was no longer recruited to the promoters of the Gata4 and Gata6 genes when R396 of Klf4 was mutated (R/K or R/A) (Figure 6I), even though these mutants still occupied the promoters themselves (FLAG ChIP, Figure 6I). These results suggested that the R396 mutation of Klf4 could not repress the PrE genes due to failed recruitment of the mSin3a complex. Therefore, we speculated that Klf4 methylation might affect its interaction with the mSin3a complex, which further regulates mSin3a complex recruitment and PrE gene expression. Co-IP assays of ectopically expressed Klf4 or its mutants in E14 cells showed that R396 mutations of Klf4 (R/K and R/A) impeded the interaction of Klf4 with the mSin3a complex (Figure 6J). Intriguingly, ChIP/re-ChIP assays confirmed that ectopic Klf4 co-occupied the promoter of Gata6 with the mSin3a complex, while the R/K mutant did not (Figure 6K). These results indicated that R396 of Klf4 is crucial for interaction with and recruitment of the mSin3a complex at the promoter regions of PrE genes. Thus, blocking the arginine methylation of Klf4 by either a R396 mutation or Prmt1 depletion resulted in decreased recruitment of the Sin3a complex to the promoter, thereby derepressing the expression of PrE genes.

To examine whether arginine methylation of Klf4 contributed to the emergence of XEN cells, XEN induction was carried out using ectopic expression of WT Klf4 and R396 mutants in E14 cells. Flow cytometry assays showed that ectopic expression of WT and R396F Klf4 markedly reduced the efficiency of XEN induction (Gata6^+^/Nanog^-^ cells), while the R396A mutant, an unmethylated form, lacked the inhibitory effects of Klf4 to some extent (Figure 6L). These results suggested that Klf4 methylation by Prmt1 was involved in preventing the emergence of XEN progenitors from mouse ES cells.

## Discussion

Arginine methylation has emerged as a prevalent posttranslational modification involved in diverse biological processes (Biggar & Li, 2015; Blanc & Richard, 2017; Guccione & Richard, 2019). Prmt1 can regulate many important processes via methylation of its histone and nonhistone substrates. The importance of Prmt1 in mouse embryo development has long been known, as Prmt1 knockout leads to early embryo lethality. However, the underlying molecular mechanisms for lethality remain unclear. In this report, we used mouse ES cells as a tool to illustrate the functions of Prmt1 in ES cell fate control. Prmt1 depletion leads to the emergence of a small population of XEN cells within ES cells, as determined using immunostaining and scRNA-Seq. Depletion of Prmt1 increases acetylation of histone H3 and H4 and leads to an open chromatin state at PrE genes. Based on careful biochemical and molecular mechanistic characterization, we suggest that Prmt1 methylates Klf4, the methylation state of which affects its binding and recruitment of the HDAC complex at PrE gene promoters for PrE gene expression. *In vivo* experiments further indicated that Prmt1 could regulate PrE lineage commitment (Figure 7).

**Figure 7.**
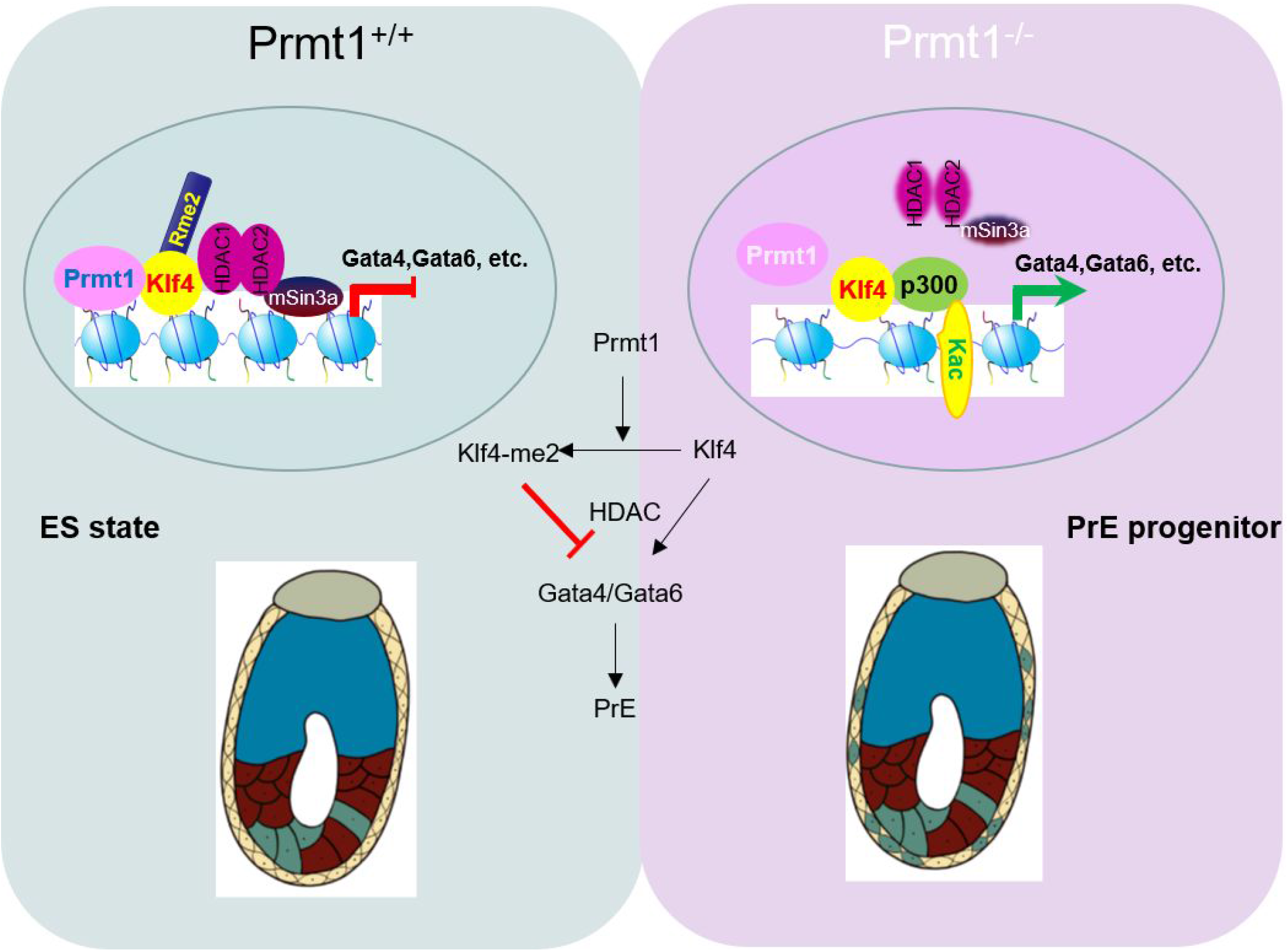
Schematic depiction of Klf4 methylation by Prmt1 to regulate PrE gene expression via mSin3a/HDAC recruitment in mouse ES cells, which heterogeneously prevents a bias toward XEN cell development.

Posttranslational modification has long been believed to be a rapid and finely tuned method of gene regulation, especially for developmental processes that call for precise modulation (Blanc and Richard, 2017). Pax7 methylation mediated by Prmt1 has been reported to regulate the muscle development process (Blanc et al., 2017). The zygotic genome needs time for reactivation after fertilization, and precise activation of genes may depend on more finely tuned switches. Posttranslational modifications of proteins and the earliest expressed regulatory factors during early embryo development seem to be good choices for better utilization of the limited regulatory switches to achieve increased precision of multilevel regulation. Relatively little is known regarding posttranslational regulation during early embryo development. Our results proved that Klf4 methylation mediated by Prmt1 may act as a regulatory factor as early as the early blastomere stage. Our study also provides a simple method to confer extraembryonic lineage commitment ability to mouse ES cells by treatment with a Prmt1 inhibitor.

Regulation of preimplantation embryo development is a fundamental biological question. Due to the short time window of preimplantation and rareness of blastomere samples, mouse ES cells are used as a tool to understand the regulatory pathways involved in mouse preimplantation development. It remains challenging to extend conclusions from ES cells into normal embryo development. The second cell fate decision, in other words, how ICM cells differentiate into PrE and EPI cells, has been an active research topic for many years. It is well known that in the ICM, a ‘salt and pepper’ expression pattern arises at E3.5 (Chazaud *et al*., 2006; Plusa *et al*., 2008). That is, some ICM cells display increased expression of PrE genes such as *Gata4* (Soudais *et al*., 1995) and *Gata6* (Fujikura *et al*., 2002; Koutsourakis *et al*., 1999; Schrode *et al*., 2014), but decreased expression of Nanog (Frankenberg *et al*., 2011) leads to sorting out and finally to the formation of the PrE layer, while the EPI fate is acquired in cells with high Nanog expression (Mitsui *et al*., 2003; Nakai-Futatsugi & Niwa, 2015; Xenopoulos *et al*., 2015). How these two portions of cells are selected remains largely unclear. The segregation of the EPI and PrE lineages in the ICM is regulated by fibroblast growth factor (FGF) signaling (Takaoka & Hamada, 2012). Fgf4, FGF receptor 2 (Fgfr2) and Grb2, which together mediate activation of the mitogen-activated protein kinase signaling pathway, are necessary for PrE formation. Our data revealed that arginine methylation of Klf4 by Prmt1 regulated the appearance of XEN cells, which provides a novel epigenetic mechanism for the second cell fate decision in mouse embryogenesis. Commitment of PrE cells can be mimicked *in vitro* by the induction of XEN cells from mouse ES cells. In the absence of Prmt1, XEN induction is highly efficient, which offers functional clues about the role of Prmt1 in PrE formation. Our results indicated that Prmt1 regulated the expression of PrE genes to bias the cell fate toward XEN cells, which could result in unbalanced cell numbers between EPI and PrE cells. These unbalanced proportions of ICM stem cells may explain why Prmt1^-/-^ mice died around the implantation of blastocysts with normal extraembryonic tissues (Pawlak *et al*., 2000). Carm1 functions in directing the initial stem cell fate *via* heterogeneous regulation of the Oct4 and Sox2 target genes (Goolam *et al*., 2016). Interestingly, our data indicated that Prmt1 might participate in the second cell fate decision in early embryo development by regulating the PrE lineage.

In addition to transcriptional activation *via* methylation of histone H4R3 (Strahl *et al*, 2001; Wang *et al*, 2001), Prmt1 participates in transcriptional regulation *via* nonhistone targets, such as Foxo1, Runx1 and C/EBPα (Liu *et al*, 2019; Yamagata *et al*, 2008; Zhao *et al*, 2008). Our data revealed that heterogeneity in the expression of PrE genes regulated by Prmt1 depends on Klf4 methylation rather than H4R3me, revealing a novel nonhistone target of Prmt1. An increasing number of studies have revealed crucial roles of the KLF in embryonic development (Bialkowska *et al*., 2017). Among the 18 members of the KLF, only Klf4 and Klf5 are expressed as early as the two-cell stage, with persistent expression until blastocyst formation (Aksoy *et al*., 2014; Azami *et al*, 2017; Blakeley *et al*, 2015; Lin *et al*, 2010), indicating their significance and similar functions. During the process of Prmt1-mediated regulation of the expression of PrE genes, Klf4 is a mediator that depends on its R396 methylation status, which gives us new insights into the role of Klf4 in ES cells and embryonic development. Notably, symmetric arginine methylation of Klf4 by Prmt5 was reported to inhibit Klf4 turnover in breast carcinogenesis (Hu *et al*, 2015), suggesting the function of Klf4 methylation. Interestingly, known genes that are heterogeneously expressed in ES cells, such as Stella (Graf & Stadtfeld, 2008; Hayashi *et al*, 2008; Sato *et al*, 2002) and Nanog (Chambers *et al*, 2003; Chambers *et al*, 2007; Silva *et al*, 2009; Singh *et al*, 2007), were downregulated in Prmt1 KO cells. All these data highlight the importance of Prmt1 in safeguarding the pluripotency of ES cells.

Combined with the scRNA-seq data, our experiments provide a microscopic view of the gene regulation process at the single-cell level. Approximately 1% of ES cells show obvious XEN features after Prmt1 knockout. This percentage can be regarded as a link between the result obtained with traditional large-scale molecular methods and microscopic single-cell data. Either Prmt1 knockout or Prmt1 inhibitor treatment predisposes mouse ES cells to XEN cell development, which should contribute to the efficient production of XEN cell lines from mouse ES cells, which are useful cell culture models of the PrE lineage (Cho *et al*, 2012; Kunath *et al*., 2005; Niakan *et al*., 2013). During chemical reprogramming, the formation of XEN-like cells marked by Gata4, Gata6, and Sox17 helps to generate iPSCs with high efficiency (Zhao *et al*, 2018; Zhao *et al*, 2015). Our data might offer new possibilities for chemical reprogramming by targeting the Prmt1/Klf4 regulatory pathway.

## Materials and Methods

### Antibodies

The antibodies used in this study were purchased from the following sources: Millipore (Billerica, MA): ASYM25 (09-814), HP1 (MAB3448), histone H3 (06-755), H3ac (06-599), H3K4me3 (07-473), H3K9me2 (07-441), H3K9me3 (07-442), H3K27me3 (07-449), H4 (05-858), H4ac (06-598), Prmt1 (07-404), and Sox2 (AB5603); Santa Cruz Biotech (Santa Cruz, CA): actin (sc-47778), c-Myc (sc-40, sc-789), GAPDH (sc-166574), Gata4 (sc-9053), Gata6 (sc-9055), GST (sc-138), mSin3a (sc-994), Oct4 (sc-5279), and p300 (sc-32244); Cell Signaling Technology (Danvers, MA): HDAC1 (2062), HDAC2 (2545s), and KLF4 (4038); Abcam (Cambridge, UK): H3R17me2 (ab412); Sigma (St. Louis, MO): FLAG (F3165); MBL (Japan): FLAG (PM020); Active Motif (Carlsbad, CA): H4R3me2a (39705); R&D Systems (Minneapolis, MN): KLF4 (AF3158); and Bethyl Laboratories (Montgomery, TX): Nanog (A300-397A).

### Plasmids and transfection

Mouse Prmt1 cDNA was amplified by PCR from P19 cells and cloned into the pcDNA6-FLAG vector. Full-length mouse KLF4 in the pCMV-Tag2B vector was a kind gift from Dr. Wen-ji Dong (PUMC). The Prmt1 mutant (E153Q) and point mutants (R to K or R to A) of Klf4 at R 387, 389, 390, 394, and 396 were generated by CR-based site-directed mutagenesis. KLF4 wild-type and mutants were cloned into the prokaryotic expression vector pGEX-4T-1 and the eukaryotic expression vector pCMV-Tag3B with a Myc-tag. All the expression plasmids above were cloned into the PyCAG-FLAG-HA vector for transfection into ESCs, and all constructs were verified by DNA sequencing.

Transfection was carried out using Lipo3000 transfection reagent (Invitrogen), according to the manufacturer’s instructions.

### Cell culture and XEN induction

Mouse E14 ES cells (a kind gift from Prof. Guo-hong Li from the Institute of Biophysics, Chinese Academy of Science) were cultured on plates coated with 0.1% gelatin (Millipore) under feeder-free conditions with 5% CO_2_ at 37°C. The ESC medium was prepared from Knockout DMEM (Gibco), with 15% fetal bovine serum (FBS, Gibco), 0.1 mM β-mercaptoethanol (Millipore), 2 mM GlutaMax (Gibco), 0.1 mM MEM nonessential amino acids (Gibco) and 1000 U/ml LIF (Millipore). HEK293T cells were cultured in DMEM supplemented with 10% FBS and maintained at 37°C with 5% CO_2_.

The ESCs was induced to develop into XEN cells following a previous protocol (Niakan *et al*., 2013). In brief, mouse ESCs were cultured in standard XEN medium: advanced RPMI 1640 (Gibco) with 15% (vol/vol) FBS (Gibco) and 0.1 mM-mercaptoethanol (Gibco). Then, the fresh cXEN derivation medium was changed to a solution of standard XEN medium supplemented with 0.01 μM all-trans retinoic acid (Sigma) dissolved in DMSO plus 10 ng/ml activin A. After induction, the cells were identified by qPCR and immunofluorescence assays.

### Chimeric assay of mouse ES cells *in vivo*

In order to detect the *in vivo* contribution of the ES cells, Ab2.2 (GFP2AtdT) cell line (Huang *et al*., 2012) was used to carry on the chimeric assay. The cells were injected into blastocysts three days after passaging, one group of which was also pretreated with Prmt1 inhibitor at 10μM for 24h before injection. For microinjection, ES cells were trypsinized by 0.05% trypsin-EDTA, and centrifuged at 1200 rpm for 3 min. The cell pellets were resuspended and replaced into a 6-well plate for 30 min to subside MEF cells. The tube containing cells was placed in the ice during the process of microinjection. Recipient blastocyst embryos were collected from super-ovulated female ICR mice at 3.5 days post-coitum (dpc). Ten digested ES cells were injected into one blastocyst, then 10-15 injected embryos were transferred to the uterus of one 2.5 dpc pseudo-pregnant ICR female. The transplanted embryos were observed at the indicated time points. The fluorescent signals of 6.5d and 12.5d embryos were detected using fluorescence stereo microscope (Leica M205FA). Besides, 6.5d embryos were fixed in 4% PFA and observed by Zeiss LSM780. All mouse experimental protocols were approved by the Institutional Animal Care and Use Committee of Peking Union Medical College & Chinese Academy of Medical Sciences. All animal care and experimental methods were carried out in accordance with the ARRIVE guidelines for animal experiments. All the mice used in this study were fed in Specific Pathogen Free (SPF) facilities.

### Immunofluorescence (IF)

The ES cells were seeded on coverslips coated with 0.1% gelatin in 6-well plates for 24 h. The cells were fixed with 4% formaldehyde for 10 min at room temperature, and the coverslips were washed two times in PBS. Then, the cells were permeabilized with 0.25% Triton-X-100 in PBS for 10 min and blocked with 1% BSA in PBS for 1 h. The cells were then incubated at room temperature with primary antibody for 1 h and with FITC-conjugated or TRITC-conjugated secondary antibody for an additional 1 h. After being washed twice in PBS and air-dried, the coverslips were mounted in anti-fade reagent with DAPI (Invitrogen). Fluorescence was detected using a Zeiss (LSM780) microscope at appropriate wavelengths.

Immunofluorescence of blastocysts for 3.5 dpc were performed as a protocol (Saiz *et al*, 2016) with some modifications. The blastocysts of 3.5dpc were collected and cultured in KSOM (Millipore, MR-020P-5F) medium in a humidified incubator under 5% CO2 at 37°C with or without 10μM Prmt1 inhibitor. After 24h, the blastocysts were rinsed in PBS-BSA (4mg/ml BSA) for three times and fixed in 4% PFA in PBS for 10 min at RT. All the steps were performed in the 35mm dish with 10mm bottom well (In Vitro Science, D35-10-1.5-N), the bottom well was coated with a layer of 1% agar in 0.9% NaCl. Fixed blastocysts were then rinsed in PBX (0.1% Triton X-100 in PBS) for 5 min and permeabilized in permeabilization solution (0.5% Triton-X, 100mM glycine in PBS) for 5 min at RT. Blastocysts were then rinsed in PBX and blocked for 30 min in the blocking solution (20% FBS with Pen-Strep in PBS). The blastocysts were then incubated in PBX containing the primary antibodies of Nanog (1:200 Invitrogen, 14-5761-80) and Gata6 (1:200 R&D, AF1700) at 4°C overnight. A layer of mineral oil was used to prevent evaporation. After blocking for 30 min, blastocysts were incubated with secondary antibodies (1:500, Donkey anti-Rabbit IgG (H+L) Highly Cross-Adsorbed Secondary Antibody, Alexa Fluor 488, Life Technology, A-21206; Donkey anti-Goat IgG (H+L) Cross-Adsorbed Secondary Antibody, Alexa Fluor 647, Invitrogen, A-21447) and Hochest 33342 (Sigma, B2261) at 4°C for 1 h. Embryos were then seeded into PBS-BSA drops in the 35mm dish and covered by mineral oil before imaging with Zeiss LSM780.

### Co-immunoprecipitation (Co-IP) and western blotting

Co-IP analyses were performed as previously described (Cheng *et al*., 2014).

### GST pulldown assay

Full-length mouse KLF4 and truncated KLF4 constructs were expressed in *E. coli* (BL21 DE3) and purified using glutathione-sepharose. GST, GST-KLF4 and truncated KLF4 were bound to glutathione-sepharose. FLAG-tagged Prmt1 was transfected into HEK293T cells, followed by incubation with bacterially expressed GST-KLF4 fusion protein sepharose in RIPA lysis buffer for 4 h at 4°C. The beads were washed 3 times using RIPA and boiled with 2× SDS loading buffer for SDS-PAGE.

### *In vitro* methylation assay and MS analysis

FLAG-Prmt1 and Prmt1m (E153Q) were expressed in HEK293T cells and immunoprecipitated with anti-FLAG M2 agarose beads (Sigma) and then eluted with excess FLAG peptides. GST-KLF4 and mutants were expressed in *E. coli* (BL21 DE3), purified using glutathione-sepharose, and eluted with glutathione. Purified Klf4 was incubated with WT Prmt1 or Prmt1m in methyltransferase buffer (50 mM Tris [pH 8.0], 1 mM PMSF, 0.5 mM DTT) along with 1 μCi of [^3^H] S-adenosylmethionine (SAM) or unlabeled 100 μM SAM at 37°C for 1 h. Reactions were stopped by adding 2× SDS-PAGE sample buffer, followed by heating at 100°C for 5 min. Samples were resolved by SDS-PAGE and stained with CBB. The destained gels were soaked in EN^3^HANCE (PerkinElmer) according to the manufacturer’s instructions and visualized by fluorography. Methylated substrates with unlabeled SAM were sent to PTM Biolabs (Hangzhou, China) for MALDI-TOF MS analysis.

### Real-time quantitative RT-PCR

Total RNA was purified with TRIzol reagent (Invitrogen) according to the manufacturer’s instructions. Two micrograms of total RNA were used for cDNA synthesis with a reverse transcription kit (Promega). Quantitative real-time PCR was performed using the SYBR premix kit (Promega), and the primer sequences used are listed in Supplemental Table S2. Gene expression was quantified by the comparative CT method and normalized to actin expression. Values are expressed as the mean ± SD. The experiments were repeated at least three times, and statistical analysis was performed on the individual experimental sets.

### RNA-Seq and analysis

Approximately 5×10^6^ E14 cells were used for each RNA-Seq experiment. The cells were washed with PBS and then lysed in 1 ml of TRIzol reagent (Invitrogen). Total RNA was extracted following the manufacturer’s instructions and dissolved in RNase-free water. A Ribo-Zero Magnetic Gold Kit (epicenter) was used to deplete ribosomal RNA within RNA samples according to the manufacturer’s protocol. The Ion Total RNA-Seq Kit v2 (Thermo Fisher Scientific) was used to prepare sequencing libraries. A total of 40~80 million reads were generated for each sample on an Ion Proton sequencing machine with an average length of 105 bp. Read quality was examined using FastQC v0.11.2. The sequencing reads were first aligned with the mouse mm10 (GRCm38.78) reference genome using RNA-STAR v2.4.0j (Dobin et al., 2013). Unaligned reads were then aligned again with Bowtie2 v2.2.4. Cufflinks v2.2.1 was used to calculate RPFMs of different genes and determine expression differences between groups. The data were visualized with IGV 2.3.40. All sequencing data were deposited in the GEO database.

### Single-cell RNA-Seq

Mouse ES cells were dissociated with TrypLE. Single ES cells were washed and diluted in PBS to a final concentration of 600 cells per μl. More than 95% cell viability was confirmed with trypan blue staining. 17.4 μl cell suspension was used to prepare single cell RNA-Seq library following ChromiumTM Single Cell 3’ Reagent Kits (Ver2) user guide. Sequencing was performed on Illumina HiSeq X ten in Rapid mode for pair end 150 bp (PE150) read length. The Seurat [https://satijalab.org/seurat/] package for cell normalization and cell filtering was applied considering the MT percentage (MT% < 20%). PCA and tSNE analysis were used to describe relationship between single cells. Graphcluster and K-mean were utilized for cell clustering and based on the marker gene achieved some cluster was combined. The Wilcox rank sum test was then used for marker gene analysis. Based on the rds file analyzed by the Seurat package, including clustering and cell marker identification, Monocle2 (Trapnell *et al*, 2014) was utilized for pseudotime analysis; the state of each cell was analyzed, and single cells were placed in clusters along a trajectory according to a biological process, such as cell differentiation by taking advantage of individual cell’s asynchronous progression of those processes. RNA velocity analysis was applied based on the mapped file of the filtered data (La Manno *et al*., 2018). The locus file was calculated, and the figure was constructed based on the Seurat clustering result.

### ChIP-qPCR assays and ChIP/re-ChIP analysis

The ChIP assays were performed as described previously (Cheng *et al*., 2014). The primers used for the qPCR assay of Gata4 and Gata6 are listed in Supplemental Table S3. The percentage of ChIP DNA relative to the input was calculated and expressed as the mean ± SD of three independent experiments.

For ChIP/re-ChIP analysis, ESCs were first transiently transfected with FLAG-KLF4 or the R/K mutant of the Klf4 expression plasmid. The sonicated cells were immunoprecipitated using anti-FLAG M2 beads (Sigma). Aliquots of the first ChIPed chromatin (1^st^ ChIP) were reverse cross-linked to obtain DNA for qPCR assays or were saved for re-IP using antibodies specific for mSin3a, HDAC1, HDAC2, or p300 for re-ChIP assays (2^nd^ ChIP). The primers are listed in Supplemental Table S3.

### DNase I sensitivity assay

The DNase I sensitivity assay was carried out as previously described (Cheng *et al*., 2014). Briefly, a total of 1× 10^7^ wild-type or Prmt1 knockout E14 cells were used. Aliquots of 10 μg DNA were purified for qPCR using the primers listed in Supplemental Table S3.

### Flow Cytometry assays

Mouse ES cells were dissociated into single cells by incubation with TrypLE (Gibco). The ells were washed and diluted in PBS. Staining was carried out using Foxp3/Transcriptn Factor Staining Buffer Set Kit (eBioscience) according to the manufactural protocol. The cells were stained with the following antibodies: anti-Nanog-APC (Cat#130-104-480, Miltenyi Biotec), anti-Gata6-PE (Cat#26452, Cell Signaling Technology), Isotype control APC (Cat#130-104-615, Miltenyi Biotec) and Isotype control PE (Cat#5742, Cell Signaling Technology). The stained cells were analyzed on a C6 flow cytometer (BD). Data were analyzed with FlowJo software.

## Acknowledgments

We thank Dr. B. Zhang of Liebing for helps in scRNA-Seq analysis. We thank Dr. Pedro Rocha of NIH for critical reading of manuscript. This work was supported by the CAMS Initiative for Innovative Medicine (2016-I2M-3-002, 2017-I2M-3-009) and the National Natural Science Foundation of China (91519301, 31871310).

## Author Contributions

Z.Z., G.Y., and Y.Z. conceived and designed the study. Z.Z., G.Y., H.W., Ya.Z., Y.C., F.C., Y. X., and M.C. performed experiments. Z.Z., G.Y., M.C., and Y.Z. analyzed the data. H.W., Y. X., Z.Z., and Y.H. performed the chimeric assay and blastocyst IF. Y.H. and Y.Z. supervised the project. Z.Z., G.Y., and Y.Z. wrote the paper. All authors discussed the results and commented on the manuscript.

